# Diversity and Structural Variability of Bacterial Microbial Communities in Rhizocompartments of Desert Leguminous Plants

**DOI:** 10.1101/2020.01.23.917765

**Authors:** Ziyuan Zhou, Guodong Ding, Minghan Yu, Guanglei Gao, Genzhu Wang

## Abstract

By assessing diversity variations of bacterial communities under different rhizocompartment types (i.e., roots, rhizosphere soil, root zone soil, and inter-shrub bulk soil), we explore the structural variability of bacterial communities in different root microenvironments under desert leguminous plant shrubs. Results will enable the influence of niche differentiation of plant roots and root soil on the structural stability of bacterial communities under three desert leguminous plant shrubs to be examined. High-throughput 16S rRNA genome sequencing was used to characterize diversity and structural differences of bacterial microbes in the rhizocompartments of three xeric leguminous plants. Results from this study confirm previous findings relating to niche differentiation in rhizocompartments under related shrubs, and they demonstrate that diversity and structural composition of bacterial communities have significant hierarchical differences across four rhizocompartment types under leguminous plant shrubs. Desert leguminous plants had significant effects on the enrichment and filtration of specific bacterial microbiomes across different rhizocompartments (*P*<0.05). The core bacterial microbiomes causing structure and composition variability of bacterial communities across different niches of desert leguminous plants are also identified. By investigating the influence of niches on the structural variability of soil bacterial communities with the differentiation of rhizocompartments under desert leguminous plant shrubs, we provide data support for the identification of dominant bacteria and future preparation of inocula, and provide a foundation for further study of the host plants-microbial interactions.

**IMPORTANCE:** Colonization by plant communities make valued contribution to sand-fixing in poor ecological desert environments, thereby reducing the effects of wind erosion in these areas. Our study revealed that specific core bacterial microbiomes in under-shrub soil microbial communities had a significant hierarchical enrichment effect among rhizocompartments, and were filtered into roots. The root endophyte microbiomes thus formed had low abundance and diversity, but their structural variability was the highest. In addition, our data also verified that the rhizocompartments of under desert leguminous plant shrubs had a significant differentiation effect for the core bacterial microbiomes enriched and filtered by host plants, and that each rhizocompartment represented a unique niche of bacterial communities. Understanding the interactions between xeric shrubs and soil microbial communities is a fundamental step for describing desert soil ecosystems, which in turn can offer a microbe-associated reference for evaluating the restoration of desert vegetation.

## INTRODUCTION

Soil microbes involved in soil carbon and nitrogen cycling exert a notable influence on global climate change (1). The effect of global climate change on desert ecosystems is noticeable (2), resulting in spatial, large-scale variations in desert vegetation communities (3). Frequent anthropogenic interferences and intensified global climate change, as well as other influencing factors, is accelerating vegetation degradation in arid, semi-arid, and dry sub-humid regions. Vegetation degradation in these areas will alter the regional ecological balance, resulting in land degradation and desertification becoming important areas of concern (4).

Vegetation restoration practices have been extensively undertaken in northwest China since the 1980s, being one of the most effective and sustainable means of controlling desertification and restoring degraded land (5). Plants such as *Hedysarum mongolicum*, *H. scoparium*, *Caragana microphylla*, and *Artemisia ordosica* are highly adaptable to arid and areas susceptible to wind-erosion, and they are believed to be suitable sand-fixing plants. In the Mu Us Desert in northwest China, these xeric shrub species are dominant plant community species (6–8), with the majority being leguminous plants.

The legume family (Fabaceae) are the third largest family of angiosperm, consisting of approximately 19,500 species distributed among 751 genera (9). Globally, leguminous plants display high degrees of species diversity and adaptability in ecosystems, including tropical rainforests, arctic tundra, alpine meadows, and the Gobi Desert (10). The nitrogen-fixing ability of leguminous plants provide an important source of biological nitrogen for agricultural and natural ecosystems, promoting sustainable agricultural production, providing necessary ecosystem services, and improving soil fertility. Many leguminous species also have important economic and ecological value, such as being sources of food, feed, materials, medicinal ingredients, and ornamental plants (11–13).

In desert environments with extreme climates, vegetation growth plays a vital role in wind prevention and sand fixation; in particular, well-developed and widely distributed plant roots help to fix and improve soil (7). Soil not only supports plant growth, it also provides nutrients required for plant growth. In turn, plants can fix external carbon sources, thereby contributing to the improvement of physical, chemical, and biological soil properties (14). Soil microbes also play a critical role in the complicated interactions between plants and soil (15, 16). It has been shown that the adaptability of desert vegetation is, to a very large extent, attributable to the influence of soil microbes (17, 18). Soil microbes participate in and control many processes of soil ecosystems, and they are important for maintaining the sustainable development and stability of soil ecosystem functions.

The plant microbiome is composed of different microbe classes, such as bacteria, archaea, oomycetes, fungi, and viruses, and it is often believed to be the second or extended genome of host plants. The microbiome not only provides functional assistance and support to host plants, it also plays a vital role in regulating the health and adaptability of individual plants and the productivity of plant communities (19–24). In turn, plant communities exert an influence on soil microbial communities through soil nutrient cycling and other ecological processes (14). For example, most soil microflorae are carbon-starved (25). Plants exude up to 40% of photosynthates to rhizospheres (26), resulting in the microbial community density of rhizosphere soil to undergo significant differential variations relative to that of bulk soil. Rhizospheres are also the areas where plants and soil microbes engage in the most intense interactions with each other.

Rhizospheres are narrow soil areas affected by root exudates, where up to 10^11^ microbial cells can be present (27) and there can be 30,000 prokaryotes per gram of root (28). Plant roots use bulk soil as a microbial diversity pool in which they can induce the enrichment of specific microbes favorable for plant growth (20). In general, species diversity of microbial communities in rhizospheres is lower than that in bulk soil (29–31). Significant differences in bacterial microbial communities between rhizosphere soil and non-rhizosphere soil is referred to as the “rhizosphere effect” (32). This phenomenon suggests that soil type is an important driver of microbial community composition in the rhizosphere (33, 34). Findings from previous studies have also indicated that core bacterial microbiomes closely associated with host plants dominate the degree of variability of the entire bacterial community across different niches (23, 35). As far as rhizocompartments are concerned as special micro-ecosystems, the relative differential variations of bacterial communities driven by various soil factors in roots, rhizosphere soil, and non-rhizosphere soil can be used to reflect the intensity of the “rhizosphere effect”, and characterize the degree of interactions between plants and soil. This method provides the opportunity to quantitatively examine bacterial microbial community diversity and structural variability in rhizocompartments of desert leguminous plants. The main hypotheses of this study, therefore, are: (i) Diversity and structural compositions of soil microbial communities present hierarchical differences across the four rhizocompartments (roots, rhizosphere soil, root zone soil, and inter-shrub bulk soil); (ii) Desert leguminous plants have a hierarchical filtering and enriching effect on the core beneficial microbes in soil via rhizocompartments, and this interaction between plants and soil microbes is one of the causes of niche differentiation among rhizocompartments under shrubs.

## RESULTS

### Alpha rarefaction curve and α-diversity of microbial communities

Results from our analysis indicate that the diversity of root endophyte microbiomes was far lower than that of under-shrub soil bacterial microbial communities. Moreover, compared to various under-shrub soil samples, root samples recorded a higher variability in the shape of rarefaction curves; root sample rarefaction curves recorded a loose distribution in the saturated amplitude range and soil sample rarefaction curves (especially rhizosphere soil samples) recorded a more concentrated distribution in the saturated amplitude range. The rarefaction curves of most root samples were generally saturated upon reaching about 1,000-1,500 OTUs (Fig. 1A), while those of under-shrub rhizosphere soil tended to be saturated upon reaching about 2,000-2,500 OTUs (Fig. 1B). The rarefaction curves of root zone soil samples and inter-shrub bulk soil samples were generally saturated when they reached about 1,500-2,500 OTUs (Fig. 1C and D). For the convenience of further statistical analyses regrading sequencing depths of various sequencing samples, we listed the Good’s coverage indices of various rhizocompartments based on more than 10,000 iterative computations in mothur (Fig. 1). Results for Good’s coverage index had a range of 95.3%-96.9%, indicating a high comparability for the sequencing depths of all rhizocompartment samples (roots, rhizosphere soil, root zone soil, and inter-shrub bulk soil). This result suggests that the sequencing depth was sufficient to reliably describe these plant rhizocompartment-related bacterial microbial communities. The Good’s coverage indices of soil samples were also significantly lower than those of root samples *(P<*0.05).

**FIG 1.**
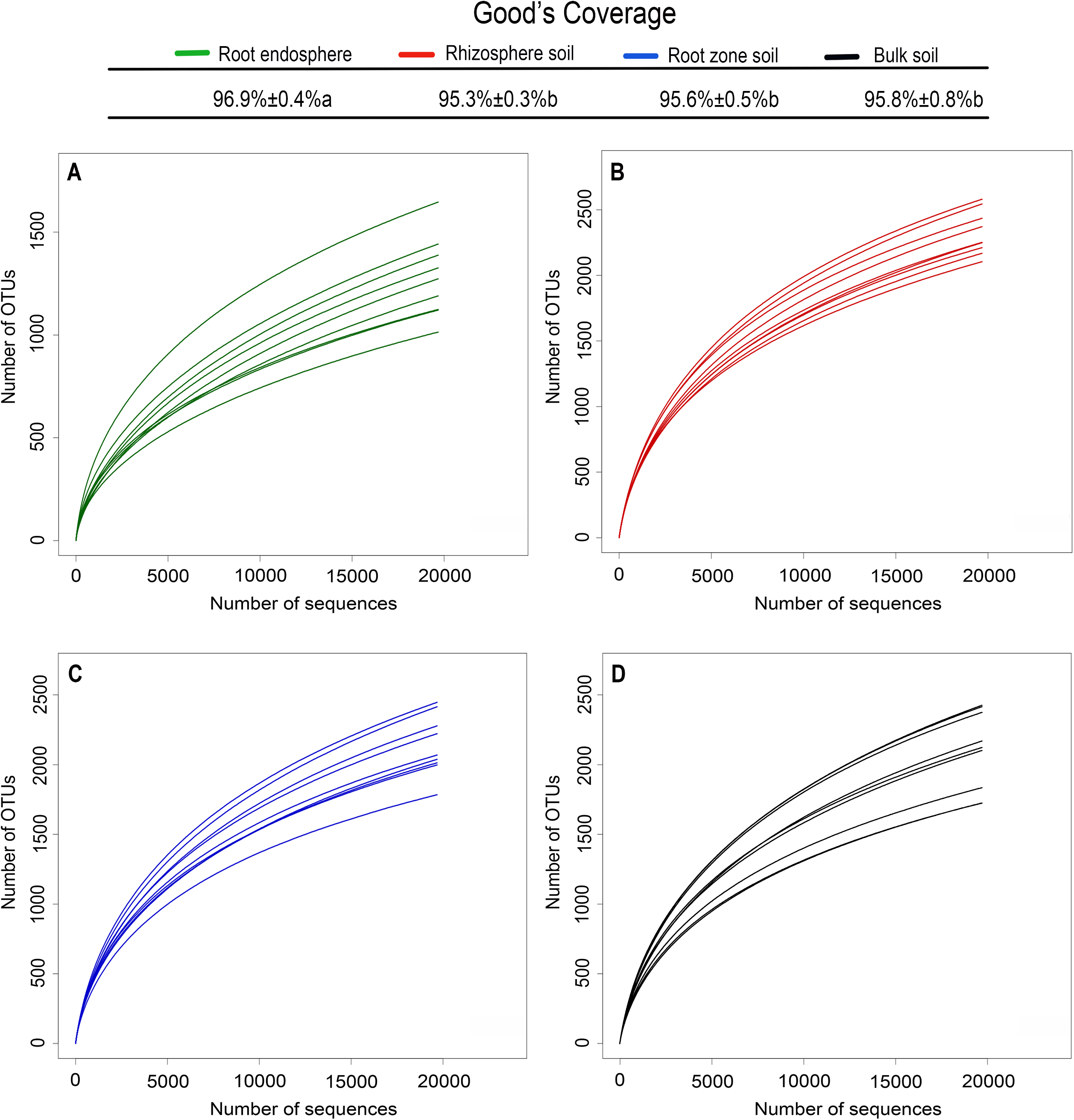
Sequencing depth estimation and rarefaction curves. (A) root samples; (B) rhizosphere soil samples; (C) root zone soil samples; (D) inter-shrub bulk soil samples. Sequencing depth estimation represents the mean ± standard deviation of nine samples of each rhizocompartment type (three plants × three replicates); lowercase letters represent statistically significant differences within the 95% confidence interval (*P*<0.05).

Based on the OTU number, Chao1 bacterial species abundance index, and Shannon microbial diversity index, α-diversity analysis was conducted on the microbial diversity of various samples (Table 1). Results indicate that an obvious separation in α-diversity existed between the root samples of the three desert leguminous plants and soil samples, that the diversity indices of under-shrub soil samples were significantly higher than those of plant root samples, and that they all attained peaks in rhizosphere soil *(P*<0.05). Specifically, OTU number, Chao1 index, and Shannon index of rhizosphere soil samples for the two *Hedysarum* L. plants were all significantly higher than those for root samples and root zone soil samples, as well as those of inter-shrub bulk soil samples *(P<*0.05). By comparison, for the three desert leguminous plants, samples of the same rhizocompartment type all showed highly similar richness and diversity estimations; and no significant effect of plant species on bacterial richness and diversity was observed (Table1).

**TABLE 1.** High-throughput genome sequencing results of the rhizocompartment samples of the three leguminous plants.

| α-diversity index | | OUT | Chao1 | Shannon |
| --- | --- | --- | --- | --- |
| Shrub species | Rhizocompartments |  |  |  |
| CM | r | 1086±63b | 1682.23±32.65b | 7.15±0.33b |
|  | rs | 2300±224a | 3244.72±306.22a | 9.50±0.19a |
|  | rz | 2260±99a | 3192.19±242.44a | 9.33±0.33a |
|  | b | 2025.57±192a | 3037.48±306.13a | 9.33±0.22a |
| HM | r | 1436±191c | 2133.36±105.03c | 7.34±0.99c |
|  | rs | 2340±148a | 3098.14±117.83a | 9.53±0.20a |
|  | rz | 2014±108b | 2790.81±100.36b | 9.08±0.05b |
|  | b | 2059±192a | 2842.44±127.15b | 9.16±0.48a |
| HS | r | 1319±126c | 2101.61±143.51c | 7.24±0.54c |
|  | rs | 2374±161a | 3309.04±130.52a | 9.56±0.16a |
|  | rz | 1998±92b | 3050.33±81.11b | 9.10±0.07b |
|  | b | 2025.07±355a | 2863.53±150.19b | 9.11±0.48a |
| $F_{\text{shrub species}}$ | | 0.57 ns | 0.14 ns | 0.10 ns |
| $F_{\text{rhizocompartments}}$ | | 47.72 | 30.64 | 71.05 |
| $F_{\text{shrub species} \times \text{rhizocompartments}}$ | | 7.70 | 8.08 | 5.47 |

Data are presented as mean ± standard deviation, n=3. CM: *Caragana microphylla*; HM: *Hedysarum mongolicum*; HS: *Hedysarum scoparium*. r: root; rs: rhizosphere soil; rz: root zone soil; b: inter-shrub bulk soil.
Different letters signify significant differences among four rhizocompartments ( $P < 0.05$ ). ns: no significant difference.
The last three rows represent the F values of the interactions between plant species/rhizocompartment types (shrub species×rhizocompartments).

### Beta diversity of microbial communities

We adopted two evolutionary phylogenetic levels (OTU and phylum) to assess the Beta diversity of microbial communities across different rhizocompartments. PCA analysis was used to display the overall similarity among various rhizocompartment samples in the structures of bacterial communities, thereby comparing the compositions of microbial communities detected in various rhizocompartments and exploring the main influencing factors driving the differences in micro-community compositions. In addition, we also adopted algorithms describing the relationships and structures of community compositions to calculate inter-sample distances, i.e., performing hierarchical clustering analysis for verification purposes based on a Bray-Curtis distance matrix (Fig. 2).

**FIG 2.**
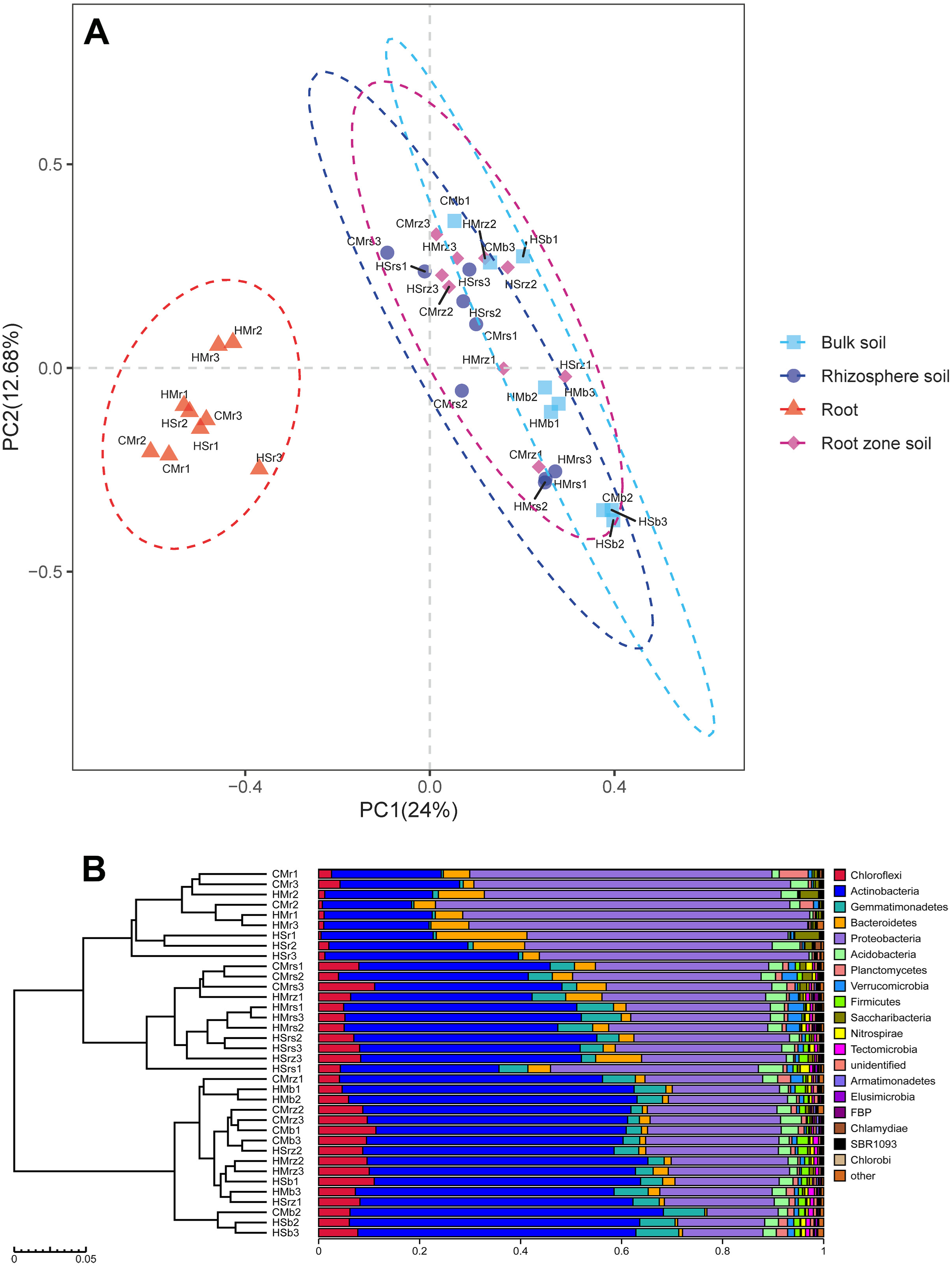
Compositions of bacterial communities driven by rhizocompartments at OTU and taxonomic levels. (A) PCA on the compositions of bacterial microbes in the rhizocompartments of desert leguminous; (B) Hierarchical clustering analysis on samples based on Bray-Curtis distance matrix. CM: *Caragana microphylla*; HM: *Hedysarum mongolicum*; HS: *Hedysarum scoparium*. r: root; rs: rhizosphere soil; rz: root zone soil; b: inter-shrub bulk soil.

PCA results (Fig. 2A) indicated that, depending on the sources of the rhizocompartments of different desert leguminous plants (roots, rhizosphere soil, root-zone soil, and inter-shrub bulk soil), bacterial communities recorded a very strong clustering performance. When PCA was based on the OTU level, PC1 and PC2 accounted for 24% and 12.68% of total variability, respectively. In addition, similar results were also obtained by grouping sources of various rhizocompartment samples and on hierarchical clustering based on a Bray–Curtis distance matrix at the phylum level, thereby verifying such clustering performance (Fig. 2B). Hierarchical clustering analysis results indicated that root samples of the three desert leguminous plants were clustered according to rhizocompartment type; the other three rhizocompartments (rhizosphere soil, root zone soil, and inter-shrub bulk soil) differed from root samples and they did not cluster completely according to their respective rhizocompartment types (Fig. 2B). To further verify the clustering performance of bacterial communities in various rhizocompartments in our PCA results, ANOSIM was performed on samples from different rhizocompartments. As indicated by analysis results, there were significant differences among various rhizocompartmental types (R=0.395, *P*=0.001) (Fig. S1).

### Analysis on of the differences in the structural compositions of major contributing bacterial taxa in various rhizocompartments

Differences in bacterial communities in the four rhizocompartments of desert leguminous plants at the phylum-order-genus levels were analyzed in-depth. At these taxonomic levels, ANOVA was used to assess the top ten ranked contributing phyla-orders-genera in terms of relative abundance percentage in the four rhizocompartments, respectively, so as to investigate the influence of rhizocompartment types on the structures and compositions of bacterial communities at various taxonomic levels (Fig. 3 and 4, and DATASET S1). In the four rhizocompartments of the three desert leguminous plants, bacterial microbes at the phylum level mainly included *Proteobacteria*, *Actinobacteria*, and *Bacteroidetes* (ten major contributing dominant bacterial phyla in total), accounting for 96%-99% of the total number of bacterial communities in the rhizocompartments (Fig. 3 and DATASET S1). The filtered major contributing bacterial phyla had significant differences in their relative abundances among the four rhizocompartments. Specifically, *Proteobacteria* presented a significant step-by-step enrichment trend in the order of root>rhizosphere soil>root zone soil (*P*<0.01) under the three desert leguminous plant shrubs; *Actinobacteria*, *Gemmatimonadetes* presented a contrary trend (*P*<0.05) (Fig. 4 and DATASET S1).

**FIG 3.**
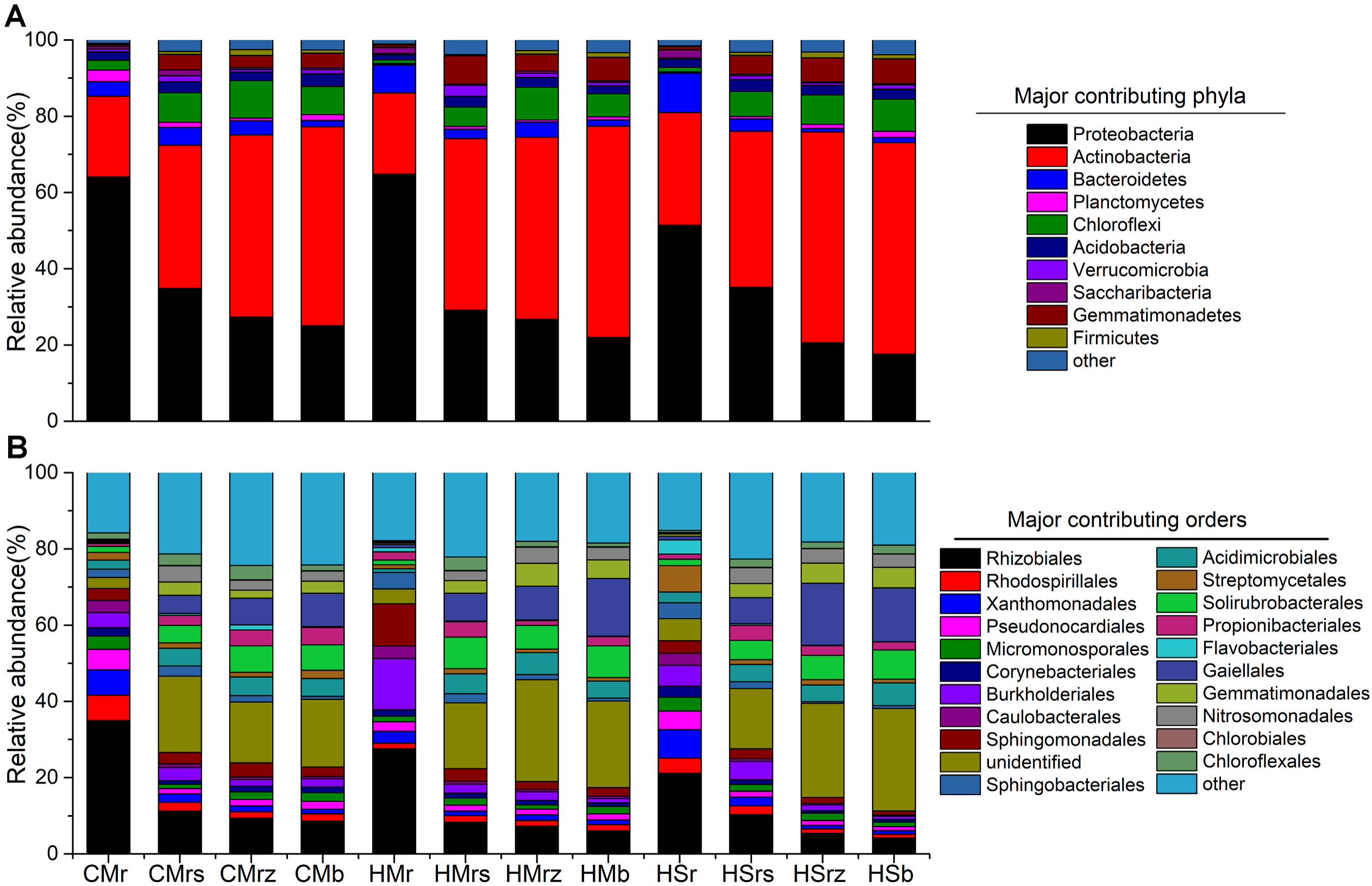
Compositions of microbial communities in the four rhizocompartments of the three leguminous plants at phylum-order levels. (A) The composition of microbial communities in the four rhizocompartments of the three leguminous plants at the phylum level; (B) The composition of microbial communities in the four rhizocompartments of the three leguminous plants at the order level. a and b list the top ten major contributing bacterial phyla and the top 20 major contributing bacterial orders, respectively, in terms of relative abundance percentage; the remaining contributors are represented by the category “other”. CM: *Caragana microphylla*; HM: *Hedysarum mongolicum*; HS: *Hedysarum scoparium*. r: root; rs: rhizosphere soil; rz: root zone soil; b: inter-shrub bulk soil. The relative abundances of major contributing bacterial taxa in the four rhizocompartments of the three leguminous plants at phylum-order-genus levels and significant effects are listed in DATASET S1.

**FIG 4.**
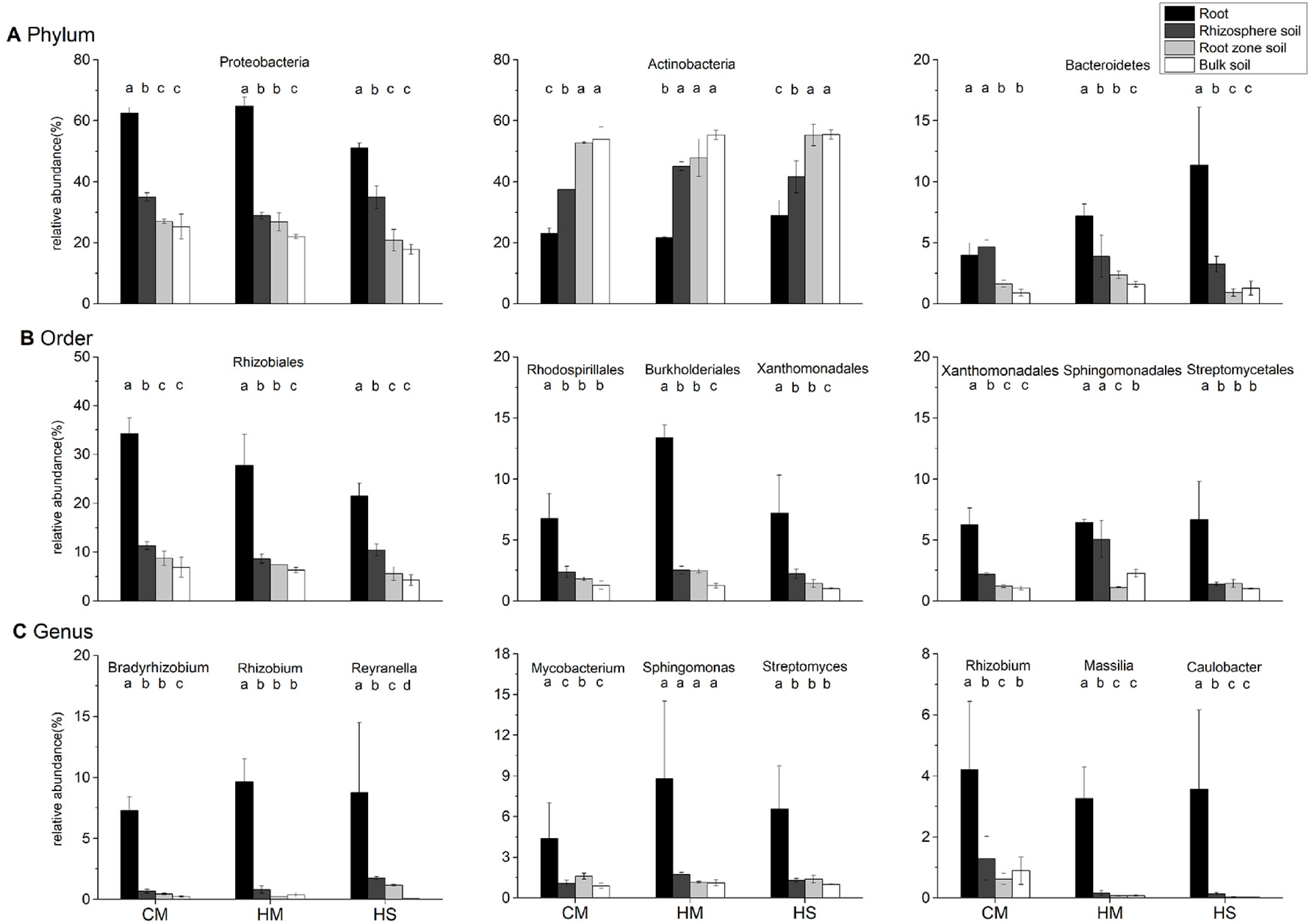
Analysis of the significance of differences in the mean relative abundances (±SE) of major bacterial communities in the four rhizocompartments of the three leguminous plants at phylum-order-genus levels. Group (A) provides the relative abundances of three major bacterial phyla; Group (B) provides the relative abundances of three major bacterial orders; Group (C) provides the relative abundances of three major bacterial genera. Different letters represent the presence of significant differences among the four rhizocompartments (*P<*0.05). Black: root; dark gray: rhizosphere soil; light gray: root zone soil; white column: inter-shrub bulk soil. Data presentation: mean ± standard deviation, *n=3*. CM: *Caragana microphylla*; HM: *Hedysarum mongolicum*; HS: *Hedysarum scoparium*.

Under the three desert leguminous plant shrubs, bacterial communities at the order level mainly included *Rhizobiales*, *Burkholderiales*, and *Sphingomonadales* (ten major contributing bacterial orders in total), accounting for 12%-75% of the total number of bacterial communities in the various rhizocompartments (Fig. 4 and DATASET S1). Specifically, *Rhizobiales* was the dominant bacterial order with the highest relative abundance under the three leguminous plant shrubs, presenting a significant trend of enrichment towards roots in all of the four rhizocompartments under leguminous plant shrubs (*P*<0.01); *Xanthomonadales* also presented a significant trend of enrichment towards roots under leguminous plant shrubs (*P*<0.05). *Burkholderiales* manifested a trend of enrichment towards roots under *H. mongolicum* shrubs (*P*<0.01), and *Rhodospirillales* showed a trend of enrichment towards roots under *C. microphylla* shrubs (*P*<0.01) and *H. scoparium* shrubs (*P*<0.02). *Bradyrhizobium*, *Rhizobium*, and *Reyranella* were the bacterial genera with the highest relative abundances in roots under *C. microphylla, H. mongolicum*, and *H. scoparium* shrubs, respectively, with all bacterial genera presenting a significant trend of enrichment towards roots (*P*<0.01). Although our results also indicated that *Rhizobium* presented an enrichment trend in *C. microphylla* roots (*P*<0.05) and *Massilia* presented an enrichment trend in *H. mongolicum* roots (*P*<0.01), some other major contributing bacterial genera also manifested a significant the same trend among the four rhizocompartments (Fig. 4 and DATASET S1).

### Enrichment and filtration effects of specific bacterial taxa among rhizocompartments

Although bacterial communities in rhizocompartments originate from under-shrub soil, our results indicate that there are varying degrees of significant differences in the structures of microbial communities among the four rhizocompartments. Therefore, in order to generate data on the microbial species that cause significant differences in microbial communities between root/rhizosphere soil/root zone soil and inter-shrub bulk soil, we used the OTU number of inter-shrub bulk soil as a control measurement. We also introduced Metastats (mothur v.1.34.4 https://www.mothur.org) to conduct significance of inter-group difference analysis on rhizocompartments and perform inter-group comparative analysis on OTUs (*P*≤0.05). Ultimately, we obtained the numbers of significantly enriched and significantly depleted OTUs of the three rhizocompartments relative to inter-shrub bulk soil. Compared to the bacterial communities of inter-shrub bulk soil, those of root/rhizosphere soil/root zone soil recorded 158/373/120 significantly enriched OTUs, and 474/97/67 significantly depleted OTUs, respectively. Based on Venn diagrams of the significantly enriched and depleted OTUs, root zone soil showed smaller differences in the structures and compositions of bacterial communities when compared to inter-shrub bulk soil (Fig. 5).

**FIG 5.**
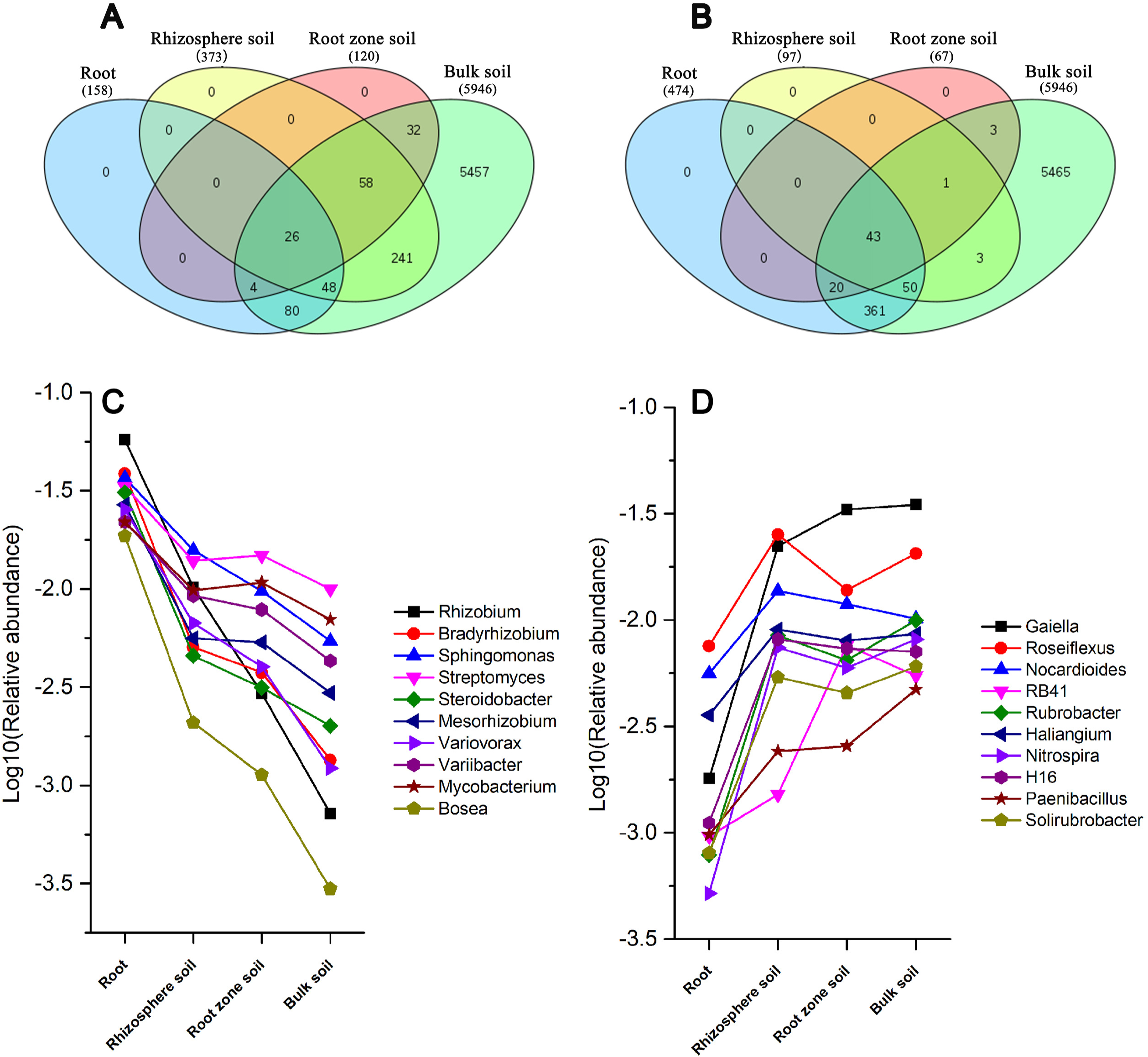
Venn diagrams of the significantly enriched (A) and depleted (B) OTUs in the other three rhizocompartments compared to root zone soil under three leguminous plant shrubs. (C) Bacterial genera corresponding to relatively enriched OTUs; (D) Bacterial genera corresponding to relatively depleted OTUs.

Enriched OTUs among various rhizocompartments all recorded varying degrees of overlapping. Specifically, among the 158 OTUs enriched in roots relative to inter-shrub bulk soil, 74/30 manifested consistent enrichment trends with OTUs from rhizosphere soil/root zone soil, respectively. Among the 373 OTUs enriched in rhizosphere soil, 84 manifested consistent enrichment trends with OTUs from root zone soil (*p*≤0.05). Twenty-six OTUs manifested an enrichment trend in all four rhizocompartment types set in this study, with rhizosphere soil recording the largest number of significantly enriched OTUs (Fig. 5A). Statistical analysis on the ten relatively abundant dominant bacterial genera corresponding to OTUs recording a relative enrichment effect among the four rhizocompartments indicated that these dominant bacterial genera all manifested a trend of enrichment towards roots (*P*≤0.05) (Fig. 5C).

Among the 474 significant relatively depleted OTUs in roots, 93/63 presented consistent depletion trends as OTUs from rhizosphere soil/root zone soil. Among the 97 relatively significantly depleted OTUs in rhizosphere soil, 44 presented consistent depletion trends as OTUs from root zone soil. Results indicate that the number of significantly depleted OTUs was small in root zone soil and highest in roots (Fig. 5B). Similarly, statistical analysis on the ten relatively abundant dominant bacterial genera corresponding to the OTUs recorded a relative depletion effect among the four rhizocompartments, indicating that these dominant bacterial genera all manifested a trend of depletion towards roots (*P*≤0.05) (Fig. 5D).

### The core bacterial microbiome in the rhizocompartments of desert leguminous plants

Results from our study indicated that rhizocompartments of desert leguminous plants had a special effect on the specific bacterial microbiome. To determine the structure and composition of this specific bacterial microbiome, we selected the top ten most abundant OTUs of each rhizocompartments as the core microbiome, resulting in a total of 24 OTUs (Fig. 6 and DATASET S2), as per the method of Beckers et al. (23). The percentages of dominant OTUs in the entire bacterial community in rhizocompartments were: 24.55% (roots), 14.54% (rhizosphere soil), 14.13% (root zone soil), and 15.70% (inter-shrub bulk soil). We then tested the effect of rhizocompartment on the normalized sequence counts of OTUs of the core community members. ANOVA analysis showed significant (*P≤*0.05) effects of plant rhizocompartments across all identified core bacterial OTUs, with the exception of *Streptomyces* (*P*=0.30), *Sphingomonas* (*P*=0.41), *Pseudarthrobacter* (*P*=0.06), *Mesorhizobium* (*P*=0.27), and *Massilia* (*P*=0.20). The core bacterial microbiome in the four rhizocompartments of the desert leguminous plants at the phylum level mainly consisted of *Proteobacteria*, *Actinobacteria*, and *Gemmatimonadetes*. *MB-A2-108* (*P*=0.005), *Gaiellales* (*P*<0.001), *Micrococcaceae* (*P*<0.001), *Gemmatimonadetes* (*P*<0.001), *Nitrosomonadaceae* (*P*<0.001) were significantly depleted in roots compared to the other three rhizocompartments. A substantial component of the core OTUs significantly enriched in roots belonged to *Bradyrhizobium* (3.74%, P<0.001), *Rhizobium* (3.28%, P<0.001), *Variovorax* (2.48%, P<0.001) and the other three genus (Fig. 6 and DATASET S2).

**FIG 6.**
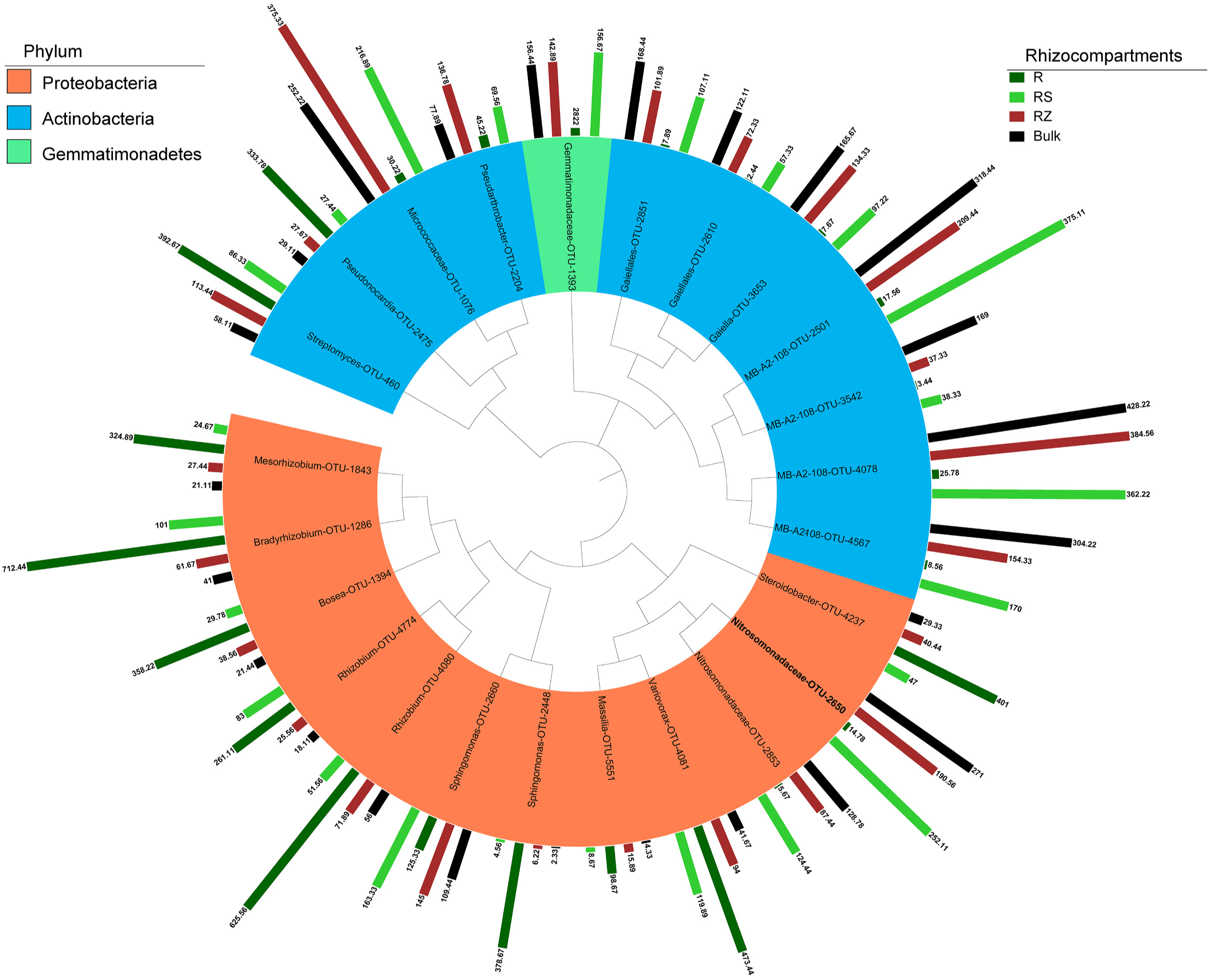
Top operational taxonomic units (OTU) members of the bacterial microbiome associated with the plant rhizocompartments. Taxonomic phylogenetic tree of core bacterial microbiomes under three desert leguminous plant shrubs. Color ranges identify phyla within the tree. Colored bars represent the relative abundance of each OTU in each plant rhizocompartments. The taxonomic dendrogram was generated with one representative sequence of each OTU using FastTree and displayed using iTOL (Interactive Tree of Life). R: root; RS: rhizosphere soil; RZ: root zone soil; Bulk: inter-shrub bulk soil. The total relative abundances of all OTUs and the significant effects across plant rhizocompartments are listed in DATASET S2.

To further explore uncovered effects potentially overlooked using core bacterial microbiome analysis, we used species indicator analyses to reveal significant associations between OTUs of the full community matrices and rhizocompartments under desert leguminous plant shrubs. This analysis identified 36 indicator OTUs in root endospheres, 53 in rhizosphere soil, 28 in root zone soil, and 146 in inter-shrub bulk soil (DATASET S3). However, when indicator OTUs of bacteria were used with an average abundance of more than 1%, eight indicator OTUs were recorded in the root samples (*P*<0.05), corresponding to *Rhizobiales*, *Gammaproteobacteria*, *Variibacter*, *Acidimicrobiales*, and *Saccharibacteria*; 17 indicator OTUs were recorded in the rhizosphere soil samples (*P*<0.05), corresponding to *Betaproteobacteria*, *Rhizobiales*, *Acidimicrobiales*, *Deltaproteobacteria*, *Gemmatimonadaceae*, *Nitrosomonadaceae*, *Bacteroidetes*, *Comamonadaceae*, *Sphingobacteriales*, *Roseiflexus*, and the *OPB35-soil-group*; 12 indicator OTUs were recorded in the root zone soil samples (*P*<0.05), corresponding to *Proteobacteria*, *Gaiellales*, *Chloroflexi*, *MB-A2-108*, *Nocardioidaceae*, *Chloroflexales*, *TK10*, *Rhodospirillales*, and *Sphingobacteriales*; and 48 indicator OTUs were recorded in the root zone soil samples (*P*<0.05), corresponding to *Actinobacteria*, *Gaiellales*, and *Solirubrobacterales*, amongst others (17 species in total) (DATASET S3).

### Relationships between rhizocompartment bacterial communities and soil factors

In this experiment, soil samples were collected from the rhizosphere, root zone and inter-shrub bulk soil of three desert leguminous plants. All soil factors of rhizosphere and root zone soils, except for NO_3_^−^-N and TP under *C. microphylla* and the two *Hedysarum spp*., recorded significant differences (*P*<0.05). Apart from a slight deficiency in NH_4_^+^-N observed in rhizosphere soils under the three shrub species, values for all other rhizosphere soil nutrient indices were uniformly higher than those in the root zone or in the inter-shrub bulk soil. In three cases, pH of the rhizosphere soil was lower than that of the root zone soil (Data of soil physicochemical factors are listed in DATASET S4).

Correlations between microbial community structure and soil physicochemical factors in rhizocompartments were calculated to identify abiotic factors that could cause variation in bacterial community diversity. Redundancy analysis (RDA) of bacterial communities in rhizocompartments revealed that the samples were divided according to soil physicochemical factors in different rhizocompartment types (Fig. 7). Among the four rhizocompartment bacterial communities, the microbiomes of root endophyte and rhizosphere soil were mainly influenced by SWC and soil nutrient; communities in the root zone soil and inter-shrub bulk soil were mainly influenced by soil pH and NH_4_^+^-N (Fig. 7A). Members of the core bacterial microbiome, which underwent hierarchical filtration and enrichment through inter-shrub bulk soil to roots by legume plants, were mainly influenced by SWC and soil nutrients. The remainder of the core bacterial microbiome, which were relatively depleted in rootscompared with the other three rhizocmpartments, were mainly influenced by soil pH and NH_4_^+^-N (Fig. 7B). Mantel test results revealed that soil pH, TN, SOC, and TP were significantly correlated with the microbial communities (*P* <0.05) (Table 2).

**FIG 7.**
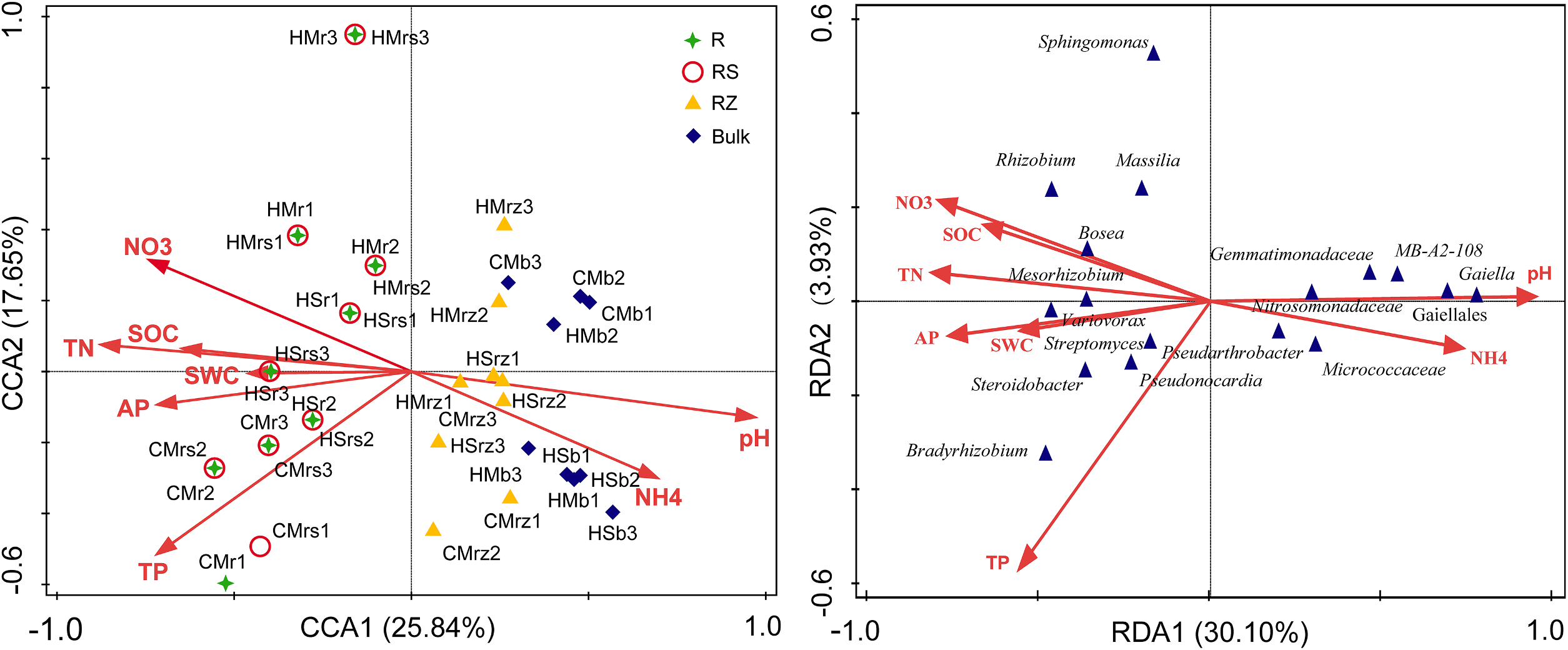
Ordination plots of the results from the redundancy analysis (RDA) on the bacterial communities in rhizocompartments and soil factors. (a) Effects of soil factors on the samples of four rhizocompartment types under three leguminous plant shrubs. (b) Effects of soil factors on the major contributing bacterial taxa (top OTU members) of the core bacterial microbiomes in four rhizocompartments under three leguminous plant shrubs. CM: *Caragana microphylla*; HM: *Hedysarum mongolicum*; HS: *Hedysarum scoparium*. r: root; rs: rhizosphere soil; rz: root zone soil; b: inter-shrub bulk soil. Data of soil physicochemical factors are listed in DATASET S4.

**TABLE 2.** Correlation between bacterial community and soil properties as shown by Mantel test.

| Soil properties | pH | TN | N-NH<sub>4</sub><sup>+</sup> | N-NO<sub>3</sub><sup>-</sup> | SOC | TP | AP | SWC |
| --- | --- | --- | --- | --- | --- | --- | --- | --- |
| *r* | 0.2478 | 0.1413 | -0.051 | -0.01 | 0.1781 | 0.1333 | 0.1299 | 0.0573 |
| *P* | **0.002** | **0.047** | 0.693 | 0.522 | **0.036** | **0.041** | 0.089 | 0.224 |

Bold type indicates significant difference ( $P < 0.05$ )

## DISCUSSION

### Structural variability of bacterial communities in rhizocompartments

When comparing root and soil samples, we observed that the rarefaction curves of OTUs recorded different saturation intervals and curve distributions upon reaching saturation. Rarefaction curve results indicate that high variability of species abundance in root endophytic OTUs of desert leguminous plants could be attributed to enrichment and filtration of specific probiotic bacteria by roots in the under-shrub soil bacterial communities under different growth states of various sample plants. A study of niches in the roots of *Populus tremula* by Beckers et al. (23) recorded similar rarefaction curve variations, indicating that uneven migration and colonization of bacterial communities with root distribution may result in these findings. Our results indicate that variations in specific core bacterial microbiomes in massive endophytes colonizing roots of various plant species constituted the primary reason for the high variability of rarefaction curves. In fact, the enrichment of bacterial communities towards rhizosphere soil mainly derived from the following sources: (i) carbon-containing primary and secondary metabolites (root exudates) in plant rhizospheres (36, 37); and (ii) allelochemicals released by plants through allelopathy, which induces chemotaxis in soil microbes (26). Although derived nutrients (such as root exudates) and chemotaxis induction exist extensively in rhizosphere niches, plant-related bacteria must undergo intense competition before successfully migrating to and colonizing rhizocompartments (38). Compared to the bacterial migration in soil among the rhizocompartments under shrubs, root endophytes must have many other properties to colonize the roots of host plants (such as the expression of chemotaxis-related genes, flagella, and the production of key enzymes for colonizing root cells) (39–41). Bacteria that enter roots must also adapt to stress factors caused by the innate immune system of host plants (42) and engage in complicated interactions with host plants to promote plant growth (40).

Results from our study indicate that, compared to soil bacterial communities under desert leguminous plant shrubs, bacterial communities in roots have a higher structural variability. It was previously highlighted that a few bacterial microbes in rhizosphere soil can enter root tissues and form endophyte microbiomes, and that their community compositions may differ from the composition of microbial communities in rhizosphere soil (43). Lundberg et al. (44) and Peiffer et al. (45) recorded that root endophyte microbiomes are not opportunistic subsets of bacterial microbial communities in rhizosphere soil, and that they are due to many complex factors, such as plant growth period, breed, and genotype. Bulgarelli et al. (46) noted that soil types determine the composition of bacterial communities in roots and, to some extent, host genotypes decide their ribotype profiles.

In summary, the colonization of endophytes in roots and the subsequent formation of a community with a relatively stable structural composition seem to constitute a highly variable process. This observation is supported by the Alpha rarefaction curve (Fig. 1), PCA and RDA results (Fig. 2 and 7), ANOVA on the relative abundances of major contributing bacterial taxa among rhizocompartments (DATASET S1), and ANOSIM on the structures of bacterial communities in this study (Fig. S1). The key to such variability lies in the colonizing capacity and properties of endophytes (40, 41), and in the fluctuations of abiotic factors (such as humidity, pH, and nutrient acquisition) among rhizocompartments (47, 48). However, OTU number, Chao1 abundance, and Shannon diversity results (Fig. 2) of rhizosphere soil, root zone soil, and bulk soil under desert leguminous plant shrubs were all higher than those of root endophyte microbiomes (Table 1).

### Drivers of the differentiation of bacterial communities under desert leguminous plant shrubs

Each rhizocompartment under the three leguminous plant shrubs were generally composed of *Proteobacteria*, *Actinobacteria*, and *Bacteroidetes* (Fig. 3, 4 and 6 and DATASET S1). This finding is consistent with the structure and composition of the dominant bacterial phyla reported in the relative abundance statistics by Sun et al. (8) on the diversity of soil microbial communities under xeric shrubs in the Mu Us Desert. The structure and composition of the major contributing bacterial phyla in root endophyte microbiomes of the three leguminous plants were basically the same. Similar to findings on root and soil bacterial communities of *Arabidopsis* (44), *Acacia* (22), *Populus* (23), soya bean, and alfalfa (49), results from our study indicate that the relative abundances of *Actinobacteria* record a significant depletion from under-shrub bulk soil to roots, and the relative abundances of *Proteobacteria* and *Bacteroidetes* presented a significant increase in the order of root zone soil<rhizosphere soil<root (*P*<0.01) and a gradual enrichment towards roots (Figure3, 4 DATASET S1). It has been shown that the trend of under-shrub enrichment of *Proteobacteria* (mostly *Alphaproteobacteria* and *Gammaproteobacteria*) towards roots is positively correlated with the available carbon pool of soil, that *Alphaproteobacteria* are closely related to heterotrophic N-fixers in high-C plots, meaning that their presence can promote an increase of NH_4_ pools (50), and that the under-shrub enrichment of *Bacteroidetes* is attributable to their ability to rapidly utilize organic matter in the soil (51). In addition, Fierer et al. (52) confirmed that the copiotrophic attributes of *Bacteroidetes* in under-shrub soil of plants were mainly contributed by *Sphingobacteriia*, and that the under-shrub enrichment of *Bacteroidetes* is not only limited by soil organic carbon content, it is also affected by other factors such as soil texture (53). However, *Actinobacteria* results in our study presented a contrary trend, recording a significant trend of depletion in the order of root<rhizosphere soil<root zone soil (*P*<0.05). Findings by Nemergut et al. (50), Van Horn et al. (54) and Lauber et al. (55) also highlighted that the relative abundance of *Actinobacteria* is positively correlated with soil pH. At the order level, *Rhizobiales* was the major contributing bacterial order with the highest relative abundance percentages in the roots of desert leguminous plants. In addition, under *H. mongolicum* shrubs, Burkholderiales showed a high degree of enrichment and significance in roots. Moulin et al. (56) and Deng et al. (57) reported that *Burkholderiales* enriched in roots may have the potential to promote plant growth. Goldfarb et al. (58) and Vandenkoornhuyse et al. (59) also highlight that *Burkholderiales* were enriched in rhizosphere soil because they can utilize root exudates to degrade aromatic compounds.

Although the degree of microbial diversity in the roots of desert leguminous plants is vast, the dominant positions of *Bradyrhizobium* and *Rhizobium* in the roots of *C. microphylla* and *H. mongolicum* are also predictable (Fig. 4 and 6, and DATASET S1) as bacteria of these two genera are known to have symbiotic rhizobia, and they are closely related to the symbiotic nitrogen fixation of leguminous plants (49, 60). However, in rhizocompartments under *H. scoparium* shrubs, neither of these two bacterial genera had prominent relative abundance percentages or enrichment significance. In *H. scoparium* roots, *Ohtaekwangia* was the bacterial genus with the highest relative abundance percentage and enrichment significance (Fig. 3 and 4, and DATASET S1). Results from previous studies examining the succession of bacterial communities in the rhizosphere soil of corn highlighted *Ohtaekwangia* to have a dominant role in the early growth phase of corn, to be readily degradable organic matter as substrates, and to be copiotrophic and fast-growing bacteria (61, 62). *Rhizobium*, *Bradyrhizobium*, and *Bosea*, as well as other dominant bacteria detected in the roots of desert leguminous plants in this study, also exist in the roots of *Acacia salicina* and *A. stenophylla* (Mimosaceae) in southeastern Australia (63, 64), and in the roots of wheat growing in volcanic ash soil in southern Chile as microsymbionts (65). The significant enrichment of the major contributing bacterial taxa under leguminous plant shrubs across the four rhizocompartments are possibly related to the hierarchical filtration of probiotic bacteria by leguminous plants (49). It has also been shown that differences between niches are possibly caused by the distribution of nutrient resources in various niches (61), and other soil abiotic factors such as soil pH, soil moisture content, and soil nutrient availability (66, 67).

Results for the mantel test identified pH, TN, SOC, and TP to be the major factors affecting bacterial community structure under desert leguminous plant shrubs (Table 2). As far as the plant species cited above are concerned, the major contributing bacterial communities in their rhizocompartments all have enriched quantities of bacteria affiliated to the Phylum *Proteobacteria* at various taxonomic levels. According to our results, *Proteobacteria* was the dominant bacterial phylum in the compositions of bacterial communities in the four rhizocompartments of the three desert leguminous plants at the phylum level. For the order level, *Rhizobiales* was the dominant bacterial order (Fig. 3-6 and DATASET S1). In the compositions of endophyte microbiomes in host plants, the vast overlapping of core microbiomes suggests that endophytic capacity (effective enriched colonization) and plant rhizocompartments (such as nutrient availability/variability. pH, and habitat suitability) are all retained for specific bacterial microbiomes (Fig. 5 and 7, and Tables S3 and S4). It is possible that the significant hierarchical enrichment and depletion trend of specific bacterial microbiomes in the roots of host plants is not just a passive process, and that it depends on the active filtration of bacterial microbiomes by host plants or the opportunistic colonization of some bacteria in suitable niches (23, 46, 68). Furthermore, interactions among plants, soil, and specific core bacterial microbiomes have led to niche differentiation in rhizocompartments.

## MATERIALS AND METHODS

### Research site

This study was undertaken at the Yanchi Research Station (37°04′-38°10′ N and 106°30′-107°47′ E) in Ningxia Province, northwest China, during September 2018. The site, located on the southern fringe of the Mu Us Desert, is characterized by a typical semi-arid continental monsoon climate that is dry and warm throughout the year. The study area has a mean annual temperature of 8.1 °C (daily temperature range: −29.4 °C ~37.5 °C; relative humidity range: 49%~55%), and a mean annual rainfall around 292 mm, 60%~70% of which falls between July and September (maximum in August). The local soil type is eolian sandy soil (69). Soil texture in the upper 1 m soil profile is classed as being sandy, having a mean bulk density of 1.5 g cm^-3^. Ecological degradation in this area is mainly caused by overgrazing, climate change, and wind erosion, resulting in the degradation of arid grasslands into sandy land. Existing vegetation was established via aerial seeding (*A. ordosica*, *H. mongolicum*, and *H. scoparium*), seedling planting (*C. microphylla*), and natural restoration (7), all of which has been undertaken since 2001. Currently, dominant xeric shrub species in this area includes *A. ordosica*, *H. mongolicum*, *H. scoparium*, *C. microphylla*, and *Salix psammophila*.

### Sampling strategy

The common dominant xeric leguminous shrub species in the study area (*C. microphylla*, *H. mongolicum*, and *H. scoparium*) were selected for analysis. These leguminous plants naturally coexist and they are widely distributed in the desert areas of northwest China (including the study area). The adaptability of these plant species to desert environments and their ecological restoration effects have previously received close attention (6–8). Samples used in this study were collected during the ripening period of test plants in September 2018. Sample plants were mainly distributed on sunny slopes of fixed dunes formed via natural restoration, and each species was sampled with three sample plots (100×100 m, with a spacing of 100 m). For each shrub, five root and soil samples were randomly collected from each sample plot. Test samples were composed of the four rhizocompartments of leguminous plants: roots, rhizosphere soil, root zone soil, and inter-shrub bulk soil. Based on local conditions, sampling methods described by Beckers et al. (70) and Xiao et al. (49) were followed when sampling the four rhizocompartment types. During sampling, plant roots were exposed and removed. Root zone soil, consisting of blocky soil by shaking and kneading from the root samples (>1 cm away from roots), was collected and stored in sterile sample bags. Soil particles adhering to the roots were collected using tweezers, was identified as rhizosphere soil to determine soil physicochemical properties. Inter-shrub bulk soil was collected at the same sampling depth as root zone soil, 10-40 cm below inter-shrub bare soil. Root samples were obtained from the secondary or tertiary branches of plant roots, and healthy and intact roots with an even thickness (5-8 cm) were removed and stored in sterile sample bags.

Root samples were initially oscillated at maximum speed for 10 min in 50 mL centrifuge tubes containing 25 mL Phosphate buffer saline (PBS; 130 mM NaCl, 7 mM Na2HPO4, 3 mM NaH2PO4, pH 7.0, 0.02% Silwet L-77). Replicate samples were rinsed with new PBS buffer with an interval of 5 min. The turbid liquid then filtered through a 100-µm nylon mesh cell strainer, and the remainder containing fine sediment and microorganisms which was the rhizosphere soil. Added 25 ml sterile PBS buffer into a new sterile 50 ml tube which contented the aforementioned root samples and vortexed, repeat this step until the PBS buffer was clear. The washed roots were placed into new centrifuge tubes containing 25 mL PBS buffer for ultrasonic oscillation for 5min; an interval of 30 s was used between the two replicates. After discarding the fluid, the clean root samples were collected. Xiao et al. (38) reported that the method for cleaning root tissues prevented contamination and damage to plant tissue samples, thereby guaranteeing the ability of treated samples to represent the rhizocompartmental properties of examined shrub species. Rhizocompartment samples from the same sample plot were mixed separately to prepare compound samples. A total of 36 DNA samples (three shrub species × four rhizocompartment types × three replicates) were prepared from the three sample plots. These samples were stored at −80 °C prior to molecular biological analysis.

In order to determine soil physicochemical properties, a conventional approach was adopted to quantify total soil organic carbon (SOC), total nitrogen (TN), total phosphorus (TP), available phosphorus (AP), ammonium nitrogen (NH_4_^+^-N), nitrate nitrogen (NO_3_^−^-N), and pH. SOC was quantified using the dichromate oxidation method (71), TN concentration of soil samples was determined using a semi-micro Kjeldahl Apparatus Nitrogen Autoanalyzer (72). AP and TP were measured using an ultraviolet spectrophotometer (UV-2550; Shimadzu, Kyoto, Japan); NH_4_^+^-N and NO_3_^−^-N were measured using the indophenol blue method and the hydrazine sulphate method, respectively; and soil pH was recorded on a 1:1 (10 g:10 mL) soil/distilled water slurry. Soil water content (SWC) was measured using a portable soil moisture meter (TRIME-PICO64/32; IMKO, Ettlingen, Germany). Due to restrictions in the observation of rhizocompartments, SWC values of rhizosphere soil and root zone soil both used observed values from the root zone in subsequent analyses.

### 16S rRNA genome sequencing and bioinformatics analysis

E.Z.N.A. soil DNA kits (OMEGA, the USA) were used with 0.5 g of fresh sample to extract DNA samples, as per the manufacturer’s instructions. All samples were stored at −80 °C. After thawing on ice, extracted DNA samples were separately centrifuged and fully mixed; sample quality was determined using a NanoDrop instrument, and 30 ng DNA was used for PCR amplification. PCR amplification was performed in 25 μL reaction volumes containing 10× PCR buffer, 0.5 μL dNTPs, 1 μL of each primer, 3 μL bovine serum albumin (2 ng/μL), 12.5 μL KAPA 2G Robust Hot Start Ready Mix, ultrapure H_2_O, and 30 ng template DNA. The PCR amplification program included initial denaturation at 95 °C for 5 min, followed by 25 cycles of denaturing at 95 °C for 45 s, annealing at 55 °C for 50 s, and extension at 72 °C for 45 s. Finally, the PCR amplification program was completed at 4 °C. Forward primer F799 (5′-AACMGGATTAGATACCCKG-3′) and reverse primer R1193 (5′-ACGTCATCCCCACCTTCC-3′) were used to target the V5-V7 regions of 16S rRNA. Both primers contained Illumina adapters, and the forward primer contained an 8 bp barcode sequence unique to each sample. An Agarose Gel DNA purification kit (Axygen Biosciences, Union City, CA, the USA) was used for the purification and combination of PCR amplicons. After purification, PCR amplicons were mixed at an equal molar concentration, followed by pair-end sequencing using the Illumina MiSeq sequencing system (Illumina, the USA) according to a standardized process.

### Statistics and analysis

MiseqPE300 (Illumina, the USA) sequencing results were recorded in the Fastq format. Quantitative insights into microbial ecology software (QIIME; Version 1.8 http://qiime.org/) was used to analyze original Fastq files and to undertake quality control according to the following criteria (73, 74): (i) base sequences with a quality score <20 at read tails were removed, and the window was set at 50 bp. When the mean quality score in the window was <20, posterior-end base sequences were discarded from the window and reads shorter than 50 bp were removed after quality control; (ii) paired reads were assembled into one sequence according to the overlapping relationship between reads (minimum overlapping length: 10 bp); (iii) the maximum allowable mismatch ratio of the overlapping areas of assembled sequences was set to 0.1, and sequences failing to meet this criterion in pairs were removed; (iv) samples were distinguished according to barcodes and primers at the head and tail ends of sequences, and the sequence directions were adjusted based on the number of mismatches allowed by barcodes (0); and (v) different reference databases were selected according to the type of sequencing data. Chimeras were removed using the Usearch program V8.1861 (http://www.drive5.com/usearch/), and clean tags of high-quality sequences were acquired after smaller-length tags were discarded using mothur software. Sequences were clustered into Operational Taxonomic Units (OTUs) using UPARSE V7.1 (http://drive5.com/uparse/) based on a 97% sequence similarity cutoff (excluding single sequences). Representative sequences and an OTU table were obtained (75).

Differences among treatments for diversity index, relative abundance data at phylum-order-genus levels and soil physicochemical factors were analyzed using one-way ANOVA model incorporating shrub species, plant rhizocompartments, and their interaction as fixed factors. Post-hoc comparison LSD tests were performed at the confidence level of 0.05. The relative abundance data of all the major contributing bacteria taxa filtered from the four rhizocompartments at phylum-order-genus levels were log-transformed, thereby adhering to the requirements for normality of data and homogeneity of variance. The Shapiro-Wilk test and the Levene test were used to test data normality and homogeneity of variance, respectively. All analyses were completed using SPSS 20.0 (SPSS Inc., Chicago, IL, USA), and α-diversity of the bacterial microbes in the rhizocompartments of desert leguminous plants was analyzed using the Vegan package (R v3.1.1). To acquire the species taxonomy information corresponding to each OTU, the RDP Classifier algorithm (http://sourceforge.net/projects/rdp-classifier/) was used on the QIIME platform (http://qiime.org/scripts/assign_taxonomy.html) to comparatively analyze the representative sequences of OTUs, and to note species information of the different communities at various levels (kingdom, phylum, class, order, family, genus, and species). Alpha rarefaction curves were constructed using OTUs with 97% sequence similarity (OTU_97_) for rarefaction analysis with Mothur V1.30.1 (http://www.mothur.org/wiki/MiSeq_SOP), and rarefaction curves were constructed using sequencing data drawn and the OTU number represented by them. The number of each sample randomly drawn was preset to start with 1, and calculated once for every increase of 50 until the rarefaction curves analyzing the OTU number of each sample all tended to be saturated on the whole. Differences in OTU composition among rhizocompartments, based on Bray-Curtis distance, were analyzed using one-way analysis of similarity (ANOSIM) with 9,999 permutations. Principal component analysis (PCA) was used to evaluate overall similarities in microbial community structure based on the Euclidean Distance. Hierarchical clustering of the samples based on Bray–Curtis dissimilarity was performed using QIIME. Mantel test was used to determine the major soil factors shaping microbial community structures. PCA, ANOSIM, and Mantel test were implemented using the Vegan package in R 3.3.1 software (76). Redundancy analysis (RDA) of the bacterial communities in the rhizocompartments was performed using CANOCO for Windows 4.5. RDA was used to analyze the relationship between microorganism and environmental factors. A taxonomic phylogenetic tree was generated with one representative sequence of each OTU using FastTree and displayed with the use of iTOL (Interactive Tree Of Life) (77).

## Acknowledgments

This study was supported by the National Key Research and Development Program of China (no. 2016YFC0500905 and 2018YFC0507102), and the National Natural Science Foundation of China (no. 31270749).

We would like to thank the staff of the Yanchi Research Station, especially Shijun Liu, Yingying He, Yuxuan Bai for their help with experimenting and sampling in the field.

The authors declare no competing financial interests.

